# Retardation of folding rates of substrate proteins in the nanocage of GroEL

**DOI:** 10.1101/2020.11.08.373423

**Authors:** Eda Koculi, D. Thirumalai

## Abstract

The *E. Coli*. ATP-consuming chaperonin machinery, a complex between GroEL and GroES, has evolved to facilitate folding of substrate proteins (SPs) that cannot do so spontaneously. A series of kinetic experiments show that the SPs are encapsulated in the GroEL/ES nano cage for a short duration. If confining the SPs in the predominantly polar cage of GroEL in order to help folding, the assisted folding rate, relative to the bulk value, should *always* be enhanced. Here, we show that this is not the case for the folding of rhodanese in the presence of the full machinery of GroEL/ES and ATP. The assisted folding rate of rhodanese decreases. Based on our finding and those reported in other studies, we suggest that the ATP-consuming chaperonin machinery has evolved to optimize the product of the folding rate and the yield of the folded SPs on the biological time scale. Neither the rate nor the yield is separately maximized.

## Introduction

The bacterial chaperonin GroEL, a stochastic nanomachine powered by ATP binding and hydrolysis ^1-2^, assists the folding of substrate proteins (SPs) that are otherwise destined for aggregation ^3-4^. GroEL consists of two heptameric rings that are stacked back-to-back ^5-6^. Each ring has seven identical subunits, which are symmetrically arranged in the resting state (absence of ligands ATP or SP), referred to as the T state ^6-8^. Like other motors ^9-10^, GroEL undergoes a series of increasingly large scale conformational (allosteric) transitions upon binding of ATP, SP, and the co-chaperonin GroES ^6-8^. In the T state the roughly cylinder shaped GroEL has a cavity ^5^. The volume of the cavity nearly doubles when both ATP and GroES are bound (referred as *R*^*I*^ or *R*^*II*^ state) ^7, 11^. The cavity volume increases roughly from 85,000Å^3^ in the T state to about 170,000Å^3^ in the *R*^*I*^ or *R*^*II*^ state ^11^. In the *R*^*II*^ state nucleotide state, the cavity can fully encapsulate a protein with a maximum of *∼* 500 amino acid residues ^12^.

How GroEL facilitates the folding of a large number of SPs that are unrelated by sequence or topology of the native state, as the machine executes complex but well-defined catalytic cycle, has remained a topic of great interest for nearly thirty years. Three scenarios for the mechanism of GroEL/ES-assisted folding have been suggested. (i) The structures of GroEL and the complex between GroEL and GroES in the ADP-hydrolyzed state shows the presence of a large cavity. From this observation it is tempting to conclude that the cavity provides the encapsulated SP a protective passive chamber in which to fold, thus avoiding aberrant interactions with other non-native SPs ^13-15^. In such an Anfinsen cage, GroEL/ES functions in a passive manner. Although theoretically the assisted folding rate when the SP is in the cage for arbitrarily long times should increase relative to the bulk value it is asserted that the rate is unchanged. (ii) In the active Anfinsen cage model, ^16-17^ it is envisioned that efficient folding in the cage occurs because kinetic traps are “entropically” disrupted, and hence the folding trajectories have unimpeded access to the folded state. There are two immediate consequences of the active cage model, which pays scant attention to the GroEL/ES catalytic cycle that occurs even in the absence of the SP albeit at a much slower rate. First, is that confinement must enhance the folding rate relative to folding in the bulk, sometimes by a factor of fifty or more ^16-17^. In contrast, theoretical arguments ^18-19^ (based on polymer theory and the ratio of the volumes of the SP to the maximum available in the cavity) and simulations have shown that the maximum acceleration in the folding rate compared to spontaneous folding cannot exceed a factor of about ten at the most. Second, the SP stability must also increase in the GroEL/ES cage. These results are rationalized using theory ^20-22^ and results from other studies ^23-26^ that confinement of the SP in the cage decreases the entropy of the unfolded state. Consequently, the folding rate must increase provided the transition state ensemble is not significantly altered. Likewise, entropy decrease of the unfolded state in the cage implies enhanced stability. The rate enhancement has been observed for certain proteins but is not universal as the active cage model asserts ^16-17, 27-29^. (iii) The Iterative Annealing Mechanism (IAM), the only theory that accounts for the coupling between the allosteric states visited by GroEL/ES during the reaction cycle and SP folding, predicts that it is the product of the folding rate and the yield of the native material (folded SP) that is maximized by repeated binding and release of the SP by GroEL/ES ^1, 12, 30-31^. Of relevance here is the implication that folding rates per se could be accelerated (modestly) or even retarded relative to the bulk. The IAM quantitatively explains the results of a large number of experiments ^12^, including observations that mutations in GroEL render it less efficient than the wild type ^12^. Recent experiments ^32-33^ have also established that in the presence of SP the chaperonin machinery responds rapidly by processing folding in both the chambers (GroEL/ES is a parallel processing machine), a discovery that accords well the IAM predictions. A consequence of the IAM is that the GroEL/ES machinery chaperones optimizes the product of the assisted folding rate (*k*_*F*_) and yield of the native material on biologically relevant time by driving the SP out of equilibrium ^34^. Neither *k*_*F*_ nor the yield is separately maximized. We note parenthetically that the same optimization principle holds for RNA chaperones.

Here, we focus on the effect of GroES/EL mediated folding rates (*k*_*F*_ s) of rhodanese, which has been extensively studied previously ^35-36^. In an important single molecule experiment, Hofmann *et al*.^29^ showed that the *k*_*F*_ of an encapsulated protein rhodanese, which is also the substrate of choice in this study), in a single ring mutant (SR1) mutant of GroEL decreases relative to the bulk due to potential interactions with the wall, as suggested using computations ^18^. Because SR1 is an artificial construct in which GroES does not disassociate from SR1 for about 300 minutes ^12^, it is unclear if a decrease in the folding rate is also observed in the full wild-type cycling system. In the full chaperonin machinery, the residence time of the SP in the expanded internal cage of GroEL is just a few seconds, and not 300 minutes ^12^. Nevertheless, we find using the cycling system consisting of GroES, GroEL, and ATP that the folding rate of rhodanese is retarded relative to its value in the bulk. Surprisingly, the extent of retardation is similar to that found in the SR1 mutant.

## Results and Discussion

To investigate the role of the wild type GroEL/ES machine on the rhodanese folding pathway, we compared the kinetics of spontaneous and assisted folding. Under our experimental conditions, we were concerned that the spontaneous folding rate (*k*_*F*_) of rhodanese released from the GroES-GroEL cage before folding is complete could contribute to the observed rate of GroEL-GroES assisted rhodanese folding. Thus, the GroEL-GroES assisted reaction was initially performed with SR1 and ADP• AlF_x_, a transition state analog of ATP. Previous experimental data have shown that GroES-SR1-ADP• AlF_x_ complex not only assisted the folding of a number of substrates, but this complex is also stable and long-lived ^37^. In the SR1 mutant, the SPs are trapped inside the GroES-SR1 ADP• AlF_x_ cage without the possibility of escaping from the cage and folding spontaneously in solution ^37^. In other words, the SPs are forced to fold in the expanded GroEL cavity.

Figure 1A shows the dependence of the extent of rhodanese folding versus time in the presence of SR1 and ATP, GroES-SR1-ADP• AlF_x_, and spontaneous folding. It is clear that, under these conditions, rhodanese is a stringent substrate, which means that there is a higher probability that this SP would fold with the assistance of the chaperonin machinery than it would otherwise. Consequently, SR1 and ATP alone are not sufficient for rhodanese folding, and the intact machinery is required to rescue the SP. Moreover, the folding of rhodanese in the presence of ATP and SR1 data demonstrate that spontaneous folding of rhodanese does not occur before the addition of GroES to the reaction mixture. The SR1 captures the unfolded rhodanese and prevents folding in solution. In other words, the pseudo first order rate for the SP capture is greater than spontaneous folding rate, which is a generic kinetic requirement for all SPs. Thus, the rhodanese folded in the presence of SR1, GroES and ADP• AlF_x_ is folded completely inside the GroES-SR1-ADP• AlF_x_ cage.

**Figure 1.**
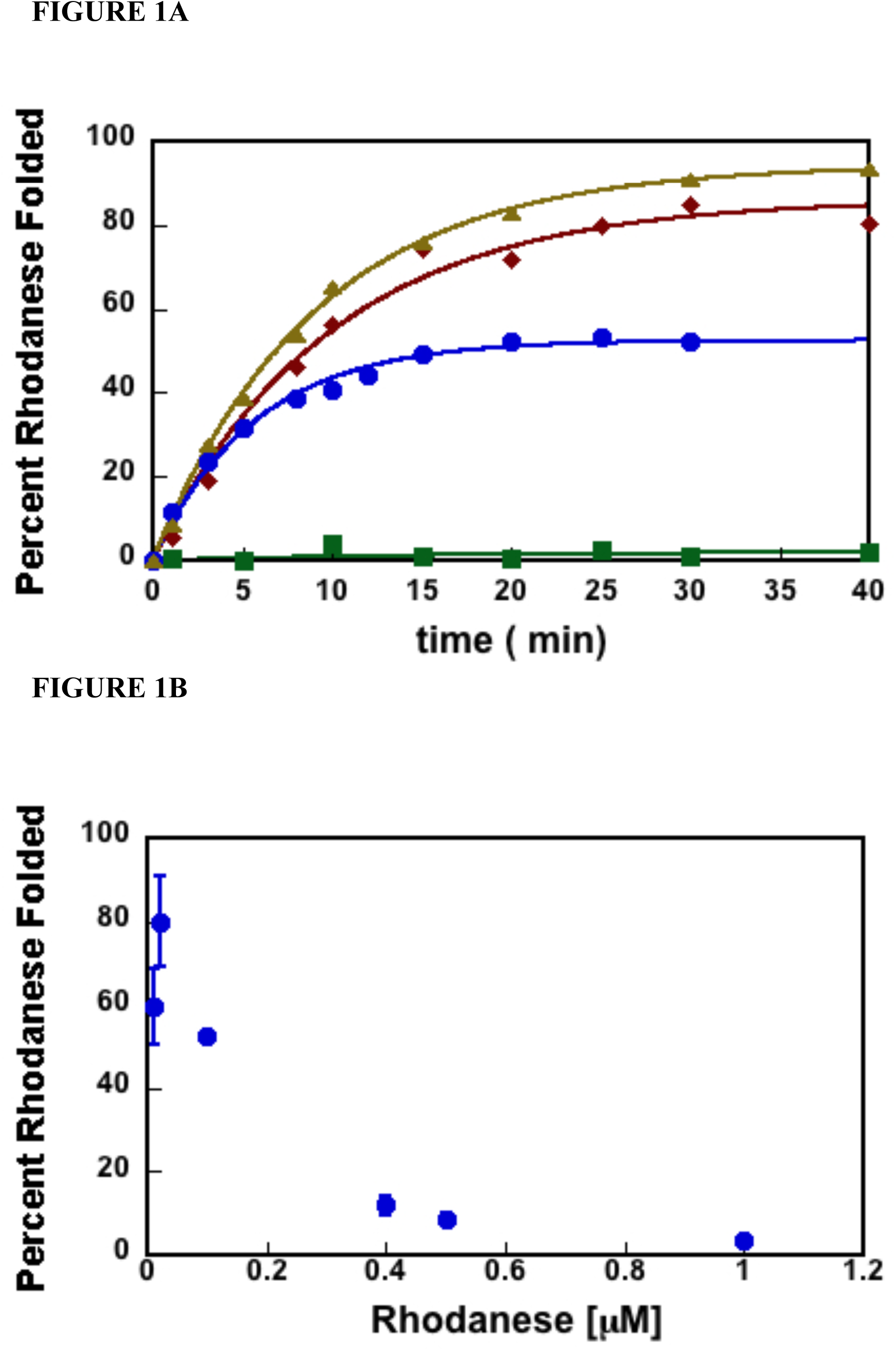
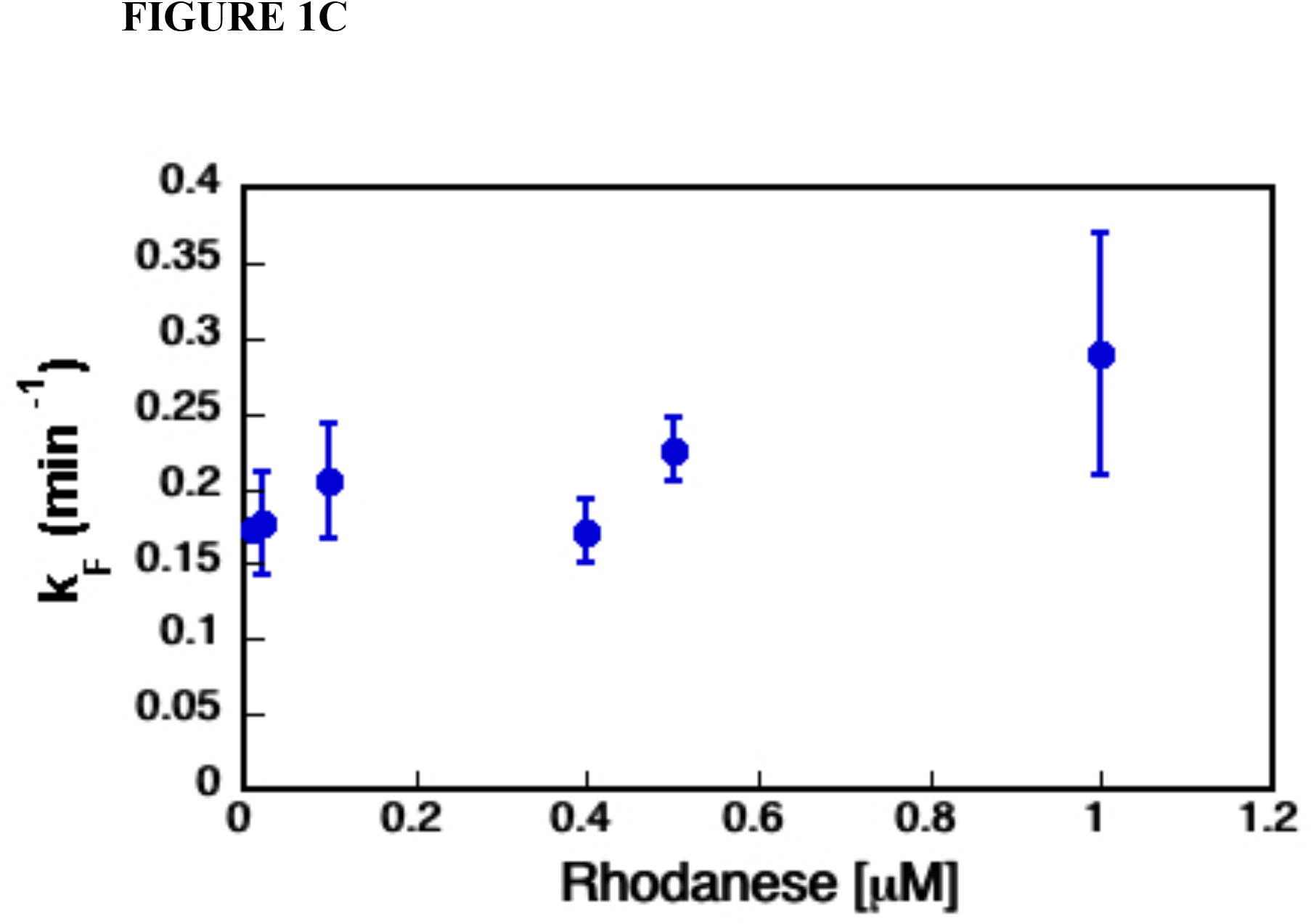
The SR1-GroES cage increases the yield of the folded SP while decreasing the SP’s rate of folding. A) Rhodanese enzymatic activity was used to monitor the extent of rhodanese folded versus time. The data shown here are representative of the folding experiments. The rate constants’ average values and standard deviations for the kinetic folding experiments are shown in Table 1. The data for rhodanese folded in presence of SR1-ADP• AlFx plus GroES is shown as brown triangles; rhodanese folded in presence of SR1 D398A-GroES-ATP plus GroES is shown as red diamonds; the rhodanese folded in presence of SR1 and ATP with no GroES as green squares; spontaneous folding of rhodanese as blue circles. B) Percentage of the rhodanese protein folded spontaneously versus the rhodanese concentration. The extent of rhodanese folded spontaneously decreases as the concentration of rhodanese increases, consistent with earlier works showing rhodanese protein is aggregation prone ^13, 43^. C) The dependence of spontaneous folding rate constant versus the rhodanese concentration. The rate of rhodanese spontaneous folding is independent of its concentration. It should be emphasized that the pseudo first order rate of conversion of the aggregated product to native rhodanese is not significant. More importantly, this process is expected to have no effect on the folding rate of native rhodanese formation from the unfolded protein. Previous work has shown that rhodanese aggregates are dead-end folding products, which do not convert to the native rhodanese over time ^44^.

More importantly, Figure 1A also shows that the yield of the folded state in the secluded SR1-GroES- ADP• AlF_x_ cage is larger than the fraction of spontaneously folded rhodanese. This is because SR1-GroES- ADP• AlF_x_ cage sequesters rhodanese for times that far exceed *k*_*F*_, thus preventing aggregation. In the bulk folding reaction, unfolded and misfolded rhodanese molecules are free to interact with each other, and form higher order aggregates. Further, evidence that spontaneously folded rhodanese has a propensity to aggregate comes from the dependence of the fraction of rhodanese folded spontaneously as a function of rhodanese concentration. As the concentration of rhodanese increases, the fraction of folded rhodanese decreases because of competing aggregation reaction that dominates as concentration of protein in solution is increased. (Figure 1B). Of particular importance here is the finding that the observed rhodanese rate of folding inside the SR1-GroES-ADP• AlF_x_ cage is two-fold slower than the spontaneous rhodanese folding rate ^29^ (Table 1).

**Table 1.** Kinetics Parameters for Spontaneous and GroEL Assisted Rhodanese Protein Folding

| | $k_F^a$<br>($\text{min}^{-1}$) |
| --- | --- |
| Spontaneous | $0.21 \pm 0.04$ |
| SR1-ADP•AlF<sub>x</sub> | $0.09 \pm 0.04$ |
| SR1 D398A | $0.1 \pm 0.01$ |
| Wild-type<br>GroEL | $0.09 \pm 0.015$ |

<sup>a</sup> Rhodanese folding rates. The values represent the average folding rate constants from at least two independent data sets and the errors are the standard deviations from these averages.

The decreased rate of folding inside the GroES-SR1-ADP• AlFx cage could be a consequence of interactions of the rhodanese with SR1-GroES cage formed in presence of ADP• AlFx, and not a consequence of rhodanese-GroEL interactions. To explore this possibility, we took advantage of the SR1 D398A construct, which forms a stable folding active chamber ^37^ when bound to GroES. The rate of rhodanese folded inside the SR1 D398A-GroES-ATP chamber, as measured by rhodanese enzyme assay, is identical within experimental errors to the rate of rhodanese folded inside the GroES-SR1-ADP•AlFx cage, and twice as slow as the rate of spontaneously folded rhodanese (Figure 1A, Table 1). Hence, the secluded SR1-GroES chamber slows down the folding process of the rhodanese protein, as reported previously ^29^.

The slower rate of folding inside the stable GroES-SR1 cage could be a consequence of the inability of the SR1 construct to progress through the allosteric states of the GroEL ATP-driven reaction cycle. To rule out this possibility, we investigated rhodanese folding using the full wild-type machinery, GroEL-GroES-ATP. Figure 2 shows the GroEL-GoES-ATP assisted rhodanese folding versus the reaction time. The observed rate of GroES-GroEL-ATP assisted rhodanese folding is very similar to the observed rates inside the SR1-GroES SR1-ADP• AlFx and SR1 D398A-GroES-ATP complexes (Table 1). Thus, the full GroEL/ES chaperonin system, while protecting the SP from aggregation, also slows down the rhodanese substrate overall rate of folding. Taken together the results of the experiments show that GroEL/ES does decrease the folding rate of the SP, contradicting the often-stated assertion that an active chaperonin always increases the rate of SP folding. Although folding in a cavity could accelerate the folding rate of SPs it is neither necessary for GroEL function nor is it valid universally.

**Figure 2.**
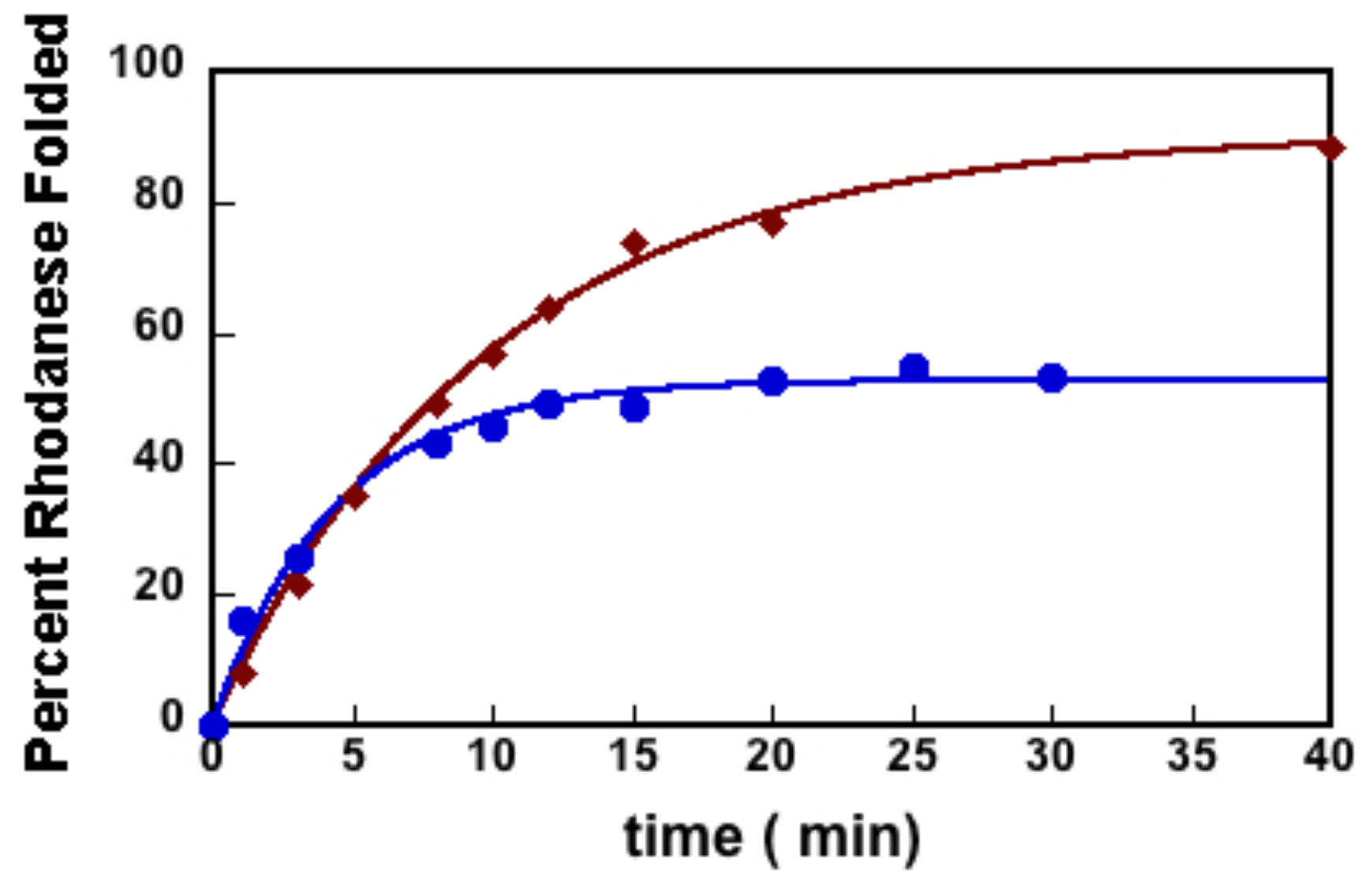
The GroEL-GroES-ATP complex increases the SP folding yield while decreases its folding rate. The extent of GroEL-ES-ATP assisted rhodanese folding versus reaction time is shown in red diamonds. The extent of spontaneous rhodanese folding versus reaction time is shown in blue circles. These data are representative of kinetic folding experiments as measured by the rhodanese enzymatic assay. The average rate constants’ values and standard deviations for the representative data depicted here are shown in Table 1.

The molecular origin in the decrease in the folding rate of the SP is hard to quantify precisely. It should be noted that the observed rate decrease, by about a factor of little over two, is not large but is clearly outside the experimental errors (see Table 1). Our finding is consistent with previous experiments ^*29*^, which reported a factor of two to about eight decrease in the rate of rhodanese folding in the SR1 mutant, depending on the temperature. One explanation, favored in the earlier work ^29^ is that the favorable interaction of the SP with exposed residues ^29^ in the interior wall of the expanded GroEL cavity ^18-19^ could increase the folding barrier. This would also imply that certain SPs are thermodynamically destabilized in the GroEL/ES cage ^38^. Regardless of the molecular mechanism, which is important to decipher, it is clear that GroEL/ES machinery has not evolved to enhance the folding rates of proteins but to maximize the yield, *Y*_*N*_, of the native material of the on biological time scales ^12-13, 29, 39^. More precisely, using theory with validation by experiments, Chakrabarty *et al*. ^34^ established that it is the product, *k*_*F*_• *Y*_*N*_, that is maximized. The chaperonin machinery has not evolved to maximize *k*_*F*_ or *Y*_*N*_ separately. In the present example, *k*_*F*_ decreases but it is compensated by an increase in *Y*_*N*_. Optimization of *k*_*F*_• *Y*_*N*_ is a general feature of protein and RNA chaperones. In the case of RNA chaperones as well, *Y*_*N*_ decreases but is compensated by an increase in *k*_*F*_ in such a way that *k*_*F*_• *Y*_*N*_ is maximized. Such an optimization is possible if chaperones drive the substrates out of equilibrium because folding of the misfolded substrate protein due to equilibrium fluctuations is only possible on time scales that far exceed biologically relevant times. Moreover, it is unlikely that GroEL/ES machine, which can processes any proteins that are unrelated by sequence or structure of the folded state, has evolved to maximize the folding rate in the cavity.

## Materials and Methods

### Protein Preparation

Bovine rhodanese bearing a C-terminal 6-His tag was purified under denaturing conditions, as previously described and stored as a lyophilized powder ^40^. GroEL, GroES, SR1, and SR1 D398A were purified as native proteins ^41^.

### Rhodanese Folding

Rhodanese was unfolded for 30 minutes at 24°C in 8 M urea, 20mM DTT and 50m Tris pH 7.5. Unfolded rhodanese was diluted to a concentration of 0.1*µ*M in the folding buffer (10 mM DTT, 50mM KCl, 10mM MgCl2, 50mM Tris pH 7.5, 50mM Na2S2O3) plus 0.2 *µ*M GroEL. The formation of rhodanese-GroEL binary complex was allowed to proceed for 5 minutes at 24ºC. Subsequently, GroES was added to the reaction mixture to a concentration of 0.4 *µ*M. The folding reaction was initiated by the addition of ATP to a final concentration of 5mM. For the SR1-ADP• AlFx assisted folding reaction, the folding buffer also contained 5mM ADP and 30mM KF, and the folding was initiated by the addition of KAl(SO4)_2_ to a final concentration of 3mM ^37^. The spontaneous rhodanese folding was initiated by diluting the unfolded rhodanese to a specific concentration in a folding buffer. The extent of correctly folded SP was measured by monitoring the absorbance at 460 nm of the complex formed between thiocyanate, one of the rhodanese reaction products, and ferric ion ^42^.

We used 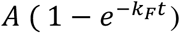, where *A* is the amplitude, *t* is the reaction time, and *k*_*F*_ is the folding rate to fit the extent of folded SP versus *t*.

## Acknowledgments

We thank Ben Schuler and Hagen Hoffmann for useful comments. This work was supported in part by NIH NRSA fellowship (F32GM079981 to EK) and Nation Institute of General Medicine (R01GM13062 to EK). DT is grateful to National Science Foundation (CHE 19-00093) and the Collie-Welch Chair (F-0019) administered through the Welch Foundation. EK performed the experiments shown in this paper in Art Horwich’s laboratory at Yale. She thanks Art Horwich and his group members for the helpful discussions.

